# Reduced-Rank Generalised Dissimilarity Modelling (RRGDM): Model-Based Ordination of Dissimilarity Data

**DOI:** 10.64898/2026.10.28.762425

**Authors:** Daniel Hernández-Carrasco, Gabrielle Koerich, Hao Ran Lai, Skipton N.C. Woolley, Jonathan D. Tonkin, Francis K.C. Hui

**Affiliations:** Faculty of Biology, University of Duisburg-Essen, Universitätsstr. 2, 45141, Essen, Germany; School of Biological Sciences, University of Canterbury, Christchurch 8140, New Zealand; Department of Aquatic Ecology, Swiss Federal Institute of Aquatic Science and Technology (Eawag), Dübendorf, Switzerland; South East Asia Rainforest Research Partnership, Kota Kinabalu, Malaysia; CSIRO Environment, Hobart, Tasmania, Australia, 7000; Te Pūnaha Matatini Centre of Research Excellence, University of Canterbury, Christchurch 8140, New Zealand; Research School of Finance, Actuarial Studies and Statistics, The Australian National University, Acton ACT 2601, Australia

## Abstract

1. Dissimilarity indices offer an effective way to represent multivariate variability in community composition and to quantify beta diversity and its drivers. However, models that regress dissimilarities against environmental predictors face the challenge of linking site-level predictors to a pairwise response.
2. This mismatch is typically circumvented by regressing dissimilarities against distances in predictor values. But this approach does not reflect how environmental variables act on individual samples, complicates the inclusion of standard modelling features such as random effects, and conflates environmental effects on average composition with effects on compositional dispersion.
3. We propose Reduced-Rank Generalised Dissimilarity Modelling (RRGDM), a framework for model-based ordination that resolves the pair-to-site mismatch by reframing dissimilarity modelling as an ordination problem. RRGDM represents dissimilarities as the outcome of sample positions in a shared, low-dimensional latent space that captures community composition. These positions are in turn modelled as a linear function of predictors and error terms, with an associated dispersion submodel.
4. Through a simulation study and two empirical examples, we show that RRGDM recovers known environmental effects and represents them as ecologically interpretable outputs. RRGDM enables inference on the relative importance of environmental gradients driving beta diversity and disentangles effects on composition versus dispersion. Although motivated by beta diversity, the approach is applicable to other pairwise dissimilarity data.

## 1 Introduction

Identifying and understanding patterns in multivariate data, such as species abundances, allele frequencies, or chemical concentrations, is a fundamental challenge across the biological sciences. The computation of pairwise dissimilarities between samples is a widely adopted solution, with a long-standing tradition (Bray & Curtis, 1957; Dice, 1945; Jaccard, 1912) that supports a versatile toolkit ranging from clustering (Everitt et al., 2011; Murtagh & Legendre, 2014) to hypothesis testing (Anderson, 2001; Anderson & Walsh, 2013) and ordination (Legendre & Anderson, 1999; Ramette, 2007). Beyond their technical utility, dissimilarities themselves are of particular interest in ecology because they can quantify beta diversity (variability in community composition), a facet of biodiversity central to community ecology and conservation (Anderson et al., 2011; Legendre & De Caceres, 2013; Socolar et al., 2016; Soininen et al., 2007). This has resulted in a distinctive way of handling dissimilarities, both as a practical means of summarising multivariate variation among communities, and as quantities of direct scientific relevance.

The study of beta diversity has motivated a large body of methods aimed at explaining community dissimilarities as a function of environmental and spatio-temporal gradients (Lichstein, 2007; Peres-Neto et al., 2006). Among model-based approaches, Generalised Dissimilarity Modelling (GDM) is a standard framework where pairwise dissimilarities are regressed against differences in predictors, often after a non-linear transformation (Ferrier et al., 2007; Fitzpatrick et al., 2025; Mokany et al., 2022). Despite their popularity, this distance-to-distance mapping limits GDM’s flexibility because many phenomena of interest are properties of samples rather than of pairs. For instance, experimental treatments and sampling designs can influence the composition of individual samples, yet representing such sample-level effects and design-based variability as pairwise differences is not straightforward (Mokany et al., 2022). Another important challenge in dissimilarity modelling is the difficulty of distinguishing changes in mean composition from changes in compositional variability (location vs. dispersion; Warton et al., 2012): samples can become more dissimilar both when their expected composition differs and when composition becomes more variable, so the two processes are often confounded at the pairwise level. This lack of separation between location and dispersion effects can complicate ecological interpretation, since the two arise from fundamentally different processes (Anderson et al., 2011). Although recent developments in the GDM literature have sought to tackle these limitations while retaining a distance-to-distance structure (Dias et al., 2022; Hernández-Carrasco et al., 2026; White et al., 2024; Woolley et al., 2017), in this article we argue that an alternative model formulation defined at the sample rather than pair level provides a more principled solution.

Ordination offers a natural template for such formulation. Because pairwise dissimilarities encode information about the relative positions of samples in a multivariate space, such as abundances of different species, ordination methods are able to recover dominant axes of compositional variation (Gower, 1966; Kruskal, 1964; Legendre & Legendre, 2012). Indeed, dissimilarity-based ordination (whether it is constrained, unconstrained, or some other variant) has proven remarkably effective in community ecology because a small number of dimensions is often sufficient to capture most of the variability among species assemblages (Allsup et al., 2023; Knight et al., 2024). To this day, these methods are routinely used to represent compositional change along environmental gradients and to identify differences in average composition and dispersion between groups of samples (e.g., Barge et al., 2019; Chase, 2010; Claar et al., 2020). However, conventional ordinations of dissimilarity matrices are descriptive rather than generative, in the sense that they do not specify a likelihood for the observed data (but see ordination methods in the context of joint species distribution models; Hui et al., 2015; van der Veen et al., 2021, 2023, among others). This means desired inferences such as uncertainty quantification and hypothesis testing are not directly available. Specifying a likelihood for dissimilarities within a model-based ordination therefore represents an opportunity to combine the strengths of dissimilarity models and conventional ordination approaches by provid-ing a sample-level formulation to the former, and a formal statistical treatment to the latter.

Here, we introduce Reduced-Rank Generalised Dissimilarity Models (RRGDM), a general framework for modelling compositional dissimilarities that bridges this gap. RRGDM treats dissimilarities as the outcome of sample positions in a low-dimensional latent space representing community composition, which is suitable for model-based ordination. Such latent composition is in turn determined by a linear predictor, making the RRGDM formulation closer to sample-level—even though dissimilarities remain the modelled response. We show that our model formulation provides enough flexibility to overcome well-known limitations of dissimilarity-based methods, including the separation of location and dispersion effects, and the inclusion of categorical predictors and random effects. Moreover, the resulting ordination space from RRGDM has attractive properties that provide an intuitive, ecologically grounded interpretation of fitted parameters, uncertainty, and model projections.

The remainder of this article is structured as follows: We first present the RRGDM framework in detail, then assess its performance in a simulation study, and demonstrate its use in two case studies. We end with a discussion of where RRGDM fits within an existing toolset for community modelling, including modern implementations of Joint Species Distribution Models, highlighting the contexts in which RRGDM may be of particular use. An R package for fitting and interrogating RRGDMs is available at (link removed for review).

## 2 Model description

Consider an *n × n* symmetric matrix ***Y***, where element *y_ij_* denotes the dissimilarity between sample *i* and *j* for *i, j* = 1*, …, n*. Also, let *x_i_* denote a set of *m* covariates (e.g., environmental factors), recorded at sample *i*. Then an RRGDM uses a known link function *g*(*·*) to connect the expected dissimilarity to the Euclidean distance between samples in a *k*-dimensional latent space *ξ*, where *k « n*:

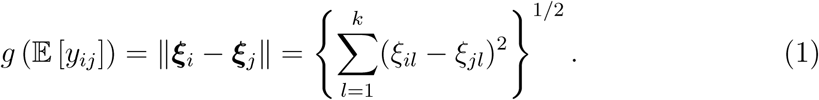

We discuss the choice of a link function and dissimilarity metric in Section 2.2. Furthermore, the position of sample *i* along dimension *l* = 1*, …, k* of the latent space is given by a linear predictor:

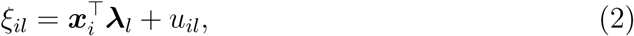

where ***λ****_l_* denotes the *m*-dimensional vector of regression coefficients quantifying the relationship between the covariates and sample positions along dimension *l*, and *u_il_*is a sample-level error term. Note no intercept term is included in ***x****_i_* such that, if the covariates are centred around their respective means, then the samples themselves will be centred at the origin of the latent space. Covariates enter (2) by positioning samples within the latent space ***ξ***; in doing so, the formulation follows the familiar structure of a linear model, thus improving interpretability and facilitating the inclusion of categorical predictors, polynomial and more general smooth terms, and random effects among other possibilities.

The sample-level error term *u_il_* represents the deviation of sample *i* from its expected position along dimension *l* according to the measured covariates, and we distinguish it from the observation-level error associated with the residual variation in dissimilarities *y_ij_* themselves discussed in the next section. Put another way, *u_il_* captures the structured residual variability whereas the observation-level pairwise error captures unstructured variability arising from, say, imperfect representation of dissimilarities in the latent space. In this article, we assume sample-level errors are independently drawn from a normal distribution *u_il_∼ N* (0*, τ* ^2^). We also assume the residual variance is common across all *k* dimensions, although this need not be the case.

Beyond modelling covariate effects on average composition, the RRGDM can be used to model compositional dispersion by additionally modelling this sample-level residual variance as a function of predictors, analogous to distributional models (Stasinopoulos et al., 2018). That is,

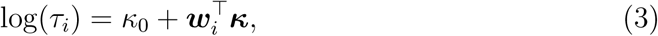

where *κ*_0_ denotes an intercept, ***w****_i_* is another set of covariate values some of the elements of which may overlap with ***x****_i_*, and ***κ*** denotes the corresponding coefficients. Equation (3) therefore quantifies how tightly or loosely sample *i* is expected to cluster around its centroid as governed by the covariates included in the dispersion sub-model. In doing so, it allows the RRGDM to simultaneously model effects on average composition (through ***λ****_l_*) versus effects on the random compositional variability (through ***κ***), separating their contribution to observed dissimilarities; see Supplementary Material for the derivation. As per (2), the model for the error variance can also include more flexible terms as appropriate (Stasinopoulos et al., 2018).

### 2.1 Response distribution

The choice of an appropriate distribution for dissimilarity data depends on the properties of the chosen function used to compute *y_ij_*. In this article, while by no means the only possibility, we focus on dissimilarity metrics that represent proportions of unshared items (e.g., species, individuals, concentrations, or biomass). These include many standard metrics in community ecology and adjacent disciplines, such as the Bray–Curtis, Sørensen–Dice, and Jaccard indices (Bray & Curtis, 1957; Dice, 1945; Jaccard, 1912). Because these are bounded between zero and one, a beta distribution serves as a possible starting choice to assume for *y_ij_* (Ferrari & Cribari-Neto, 2004). On the other hand, dissimilarities are often ratios of counts and are therefore not strictly continuous. Taking the Sørensen–Dice dissimilarity as an example, *y_ij_* = (*b_i_* + *b_j_*)*/*(2*a_ij_*+ *b_i_* + *b_j_*), where *a_ij_* represents the number of items shared between samples *i* and *j* and *b_i_* & *b_j_* represent the number of items unique to samples *i* and *j* respectively, it is clear that if *a_ij_, b_i_*, and *b_j_* are all integers then the distribution of *y_ij_* will be discrete in most realistic applications. With this in mind, we follow the work of Hernández-Carrasco et al. (2026) and Woolley et al. (2017) and assume a binomial distribution for *y_ij_*in the RRGDM, where the numerator and denominator in the formula above correspond to the number of ‘successes’ and number of ‘trials’, respectively:

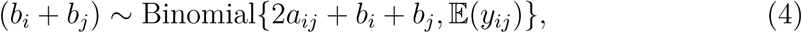

where the mean dissimilarity follows (1). If additional overdispersion is required, then the binomial distribution can be replaced by a beta-binomial distribution (Griffiths, 1973), which preserves the discrete nature of the binomial distribution while allowing greater flexibility for residual overdispersion in the response.

### 2.2 Ordination, link function, and dissimilarity metrics

RRGDM offers a means of model-based ordination based on dissimilarities. Through (1), the expected dissimilarity between two samples depends only on their Euclidean distance in the latent *k*-dimensional space (i.e., the latent space is isotropic). This facilitates the interpretation of the ordination: distances between samples can be compared directly, and the axes carry no further meaning beyond acting as a coordinate system. It also implies that the dispersion of samples across different regions of the ordination remains comparable.

Further ecological interpretation of RRGDM parameters and ordination depends on the choice of a dissimilarity metric and link function. In the two case-studies in Section 5, we favour a combination that deliberately makes distances in the latent space interpretable as constant rates of compositional change. Specifically, we combine dissimilarity metrics that represent proportions of unshared items (e.g., Sørensen–Dice and Bray–Curtis dissimilarities) with the so-called negative log link, 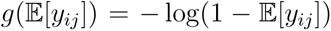. Equivalently, from (1) we have the inverse link relationship 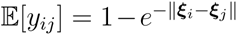 (Fig. 1). Note with this link choice, similarity declines exponentially with distance in the latent space, i.e., 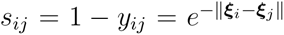, at a constant rate *d* ln (*s*_*ij*_)/*d* ||***ξ****_i_ − **ξ**_j_||* = -1) (Nekola & White, 1999), such that each unit increase in latent distance corresponds to the same proportional reduction in similarity.

**Figure 1:**
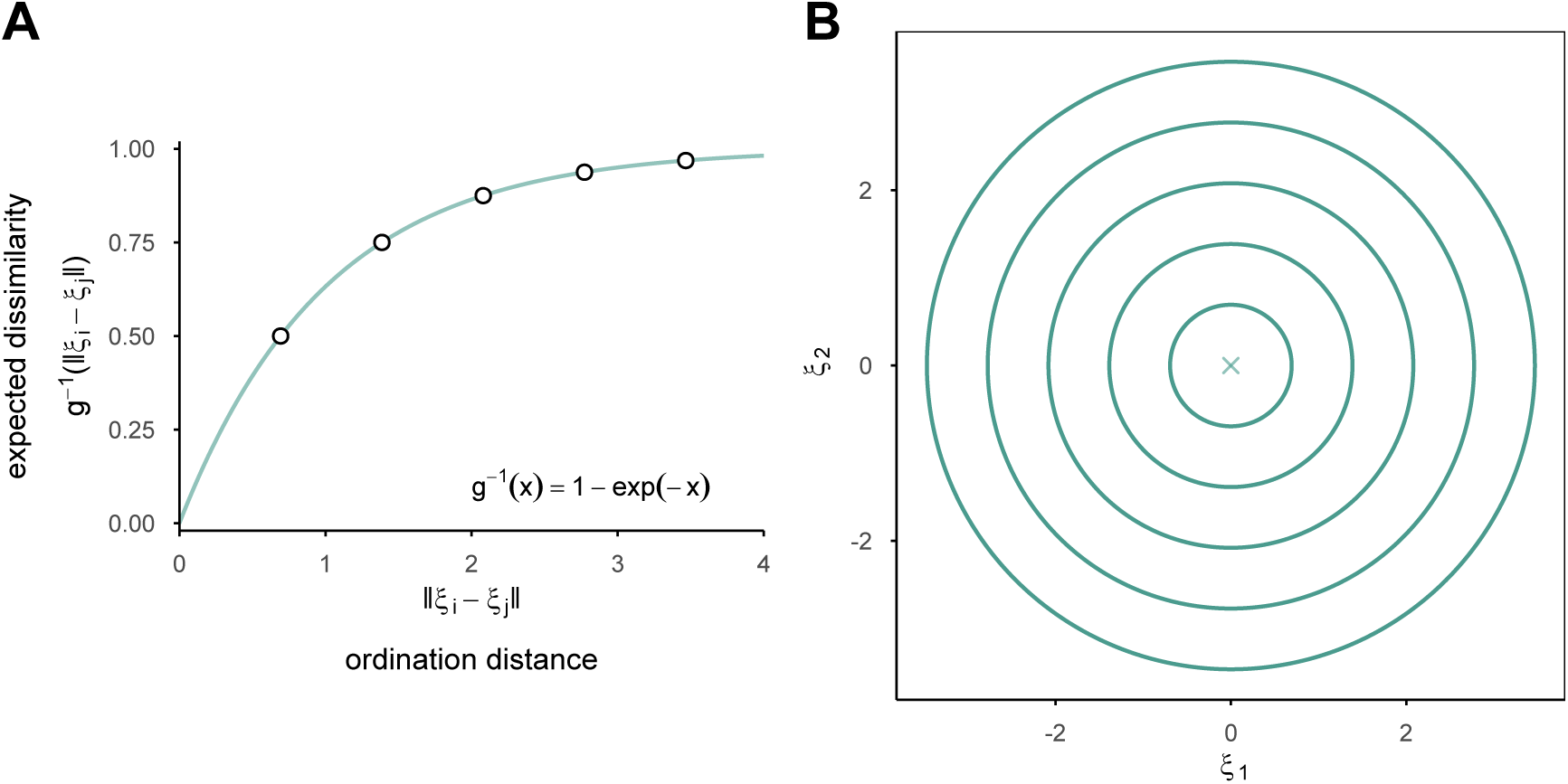
Relationship between pairwise distances in RRGDM latent space and expected dissimilarities. Under the negative log link function, equal steps in latent distance correspond to equal proportional losses of similarity, so distances in the ordination can be read directly as rates of compositional turnover. (A) Shape of the negative log link function. Circles mark dissimilarity values given by *y* = 1 *−* 0.5*^n^* with *n* = 1, 2*, …,* 5, corresponding to successive 50% decreases in similarity. (B) Mapping of dissimilarities as distances in the RRGDM ordination, with *k* = 2. The concentric circles represent contours of equal distance from the origin (marked with cross), where the distances are again based on *y* = 1 *−* 0.5*^n^* with *n* = 1, 2*, …,* 5.

To see how this property improves the interpretability of RRGDM ordination, consider the Sørensen-Dice dissimilarity again, but where the two samples of communities share half of their species. This corresponds to a latent distance of ||***ξ****_i_ − **ξ**_j_||* = ln(2). Doubling their distance to ||***ξ****_i_ − **ξ**_j_||* = 2 ln(2) implies halving the proportion of shared species, increasing the expected dissimilarity from 0.50 to 0.75. Increasing the distance again to ||***ξ****_i_ − **ξ**_j_||* = 3 ln(2) halves the remaining proportion of shared species and increases dissimilarity from 0.75 to 0.875. Such a property makes the negative log link a general and robust default, and is consistent with the broad empirical observation that log-similarity decays approximately linearly with environmental and spatial distance (Nekola & White, 1999; Soininen et al., 2007).

In addition to the link function, the adequacy of RRGDM partly depends on its dimensionality *k*. Predictors can explain at most the variability captured by those *k* dimensions, so how well the latent space reproduces the observed dissimilarities sets an upper bound on explanatory power. This can be quantified using a stress metric such as:

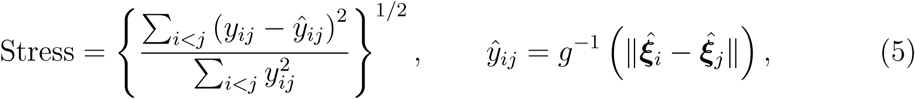

where 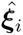 are the conditional latent coordinates (i.e., including the sample-level errors *u_il_*). This stress metric (5) is similar to other functions used in multidimensional scaling (de Leeuw & Mair, 2009; Kruskal, 1964), but intentionally involves neither a rank-based transformation of the dissimilarities nor a rescaling of the configuration, because the link function in RRGDM already maps latent distances onto the scale of dissimilarities. A stress of 0.2 or higher, conventionally taken as indicative of poor representation, implies that the root mean square error of the fitted dissimilarities is greater than 20% of the root mean square of the observed ones.

## 3 Estimation and inference

Let **Λ** = [***λ***_1_ ***λ***_2_ … λ_k_] denote the *m × k* matrix of covariate effects, and ***U*** denote the *n × k* matrix of sample-level errors where the (*i, l*)th element is given by *u_il_*. Then it is straightforward to show that, without additional constraints, the parameters in the RRGDM are not identifiable because orthogonal transformations (e.g., rotation and reflections) of **Λ** and ***U*** leave pairwise distances in the latent space unchanged. To overcome this, we use the common solution of imposing a lower triangular structure on **Λ** along with a positive constraint on its diagonal elements (Hui et al., 2015; van der Veen et al., 2023) i.e., *λ_rs_* = 0 for all *s > r* and *λ_rr_ >* 0 for *r* = 1*, …, k*. We note that full identifiability under this constraint requires *m ≥ k*, but rotational invariance is already removed when *m ≥ k −* 1. The latter yields two mirror-image optima rather than a continuous ridge, which is far less problematic for optimisation and inference. When the conditions for identifiability cannot be met, analogous constraints on the matrix of sample-level errors ***U*** are also possible although achieving convergence can be more difficult given their stochastic nature.

For all simulations and case studies herein, we fit RRGDMs by maximum likelihood estimation via Template Model Builder (TMB, Kristensen et al., 2016), applying the Laplace approximation to integrate out the sample-level errors ***U*** (along with any other random effects) and obtain the approximate marginal log-likelihood. As the optimisation routine, we used the nlminb function in R, while standard errors are obtained from TMB’s sdreport function. To further reduce computation time and assist in model convergence, starting values are obtained from a Principal Co-ordinates Analysis of the link-transformed dissimilarities followed by distance-based Redundancy Analysis, rotated to satisfy the identifiability constraint (Supplementary Material). Starting values may also be jittered for multiple restarts (analogous to Niku et al., 2019).

After fitting, point predictions of ***ξ****_i_* and associated derived quantities are directly obtained from TMB, while associated uncertainty intervals for these can be obtained by simulating parameter draws from the asymptotic multivariate normal distribution of the estimates (single-fit bootstrapping, Fletcher & Jowett, 2022). Next, the relative importance of a specific covariate, say *x_ip_*, in driving directional changes in composition (location effect) can be summarised by the modulus of its corresponding coefficient vector across the *k* latent dimensions i.e., ||***λ****_p,·_*|| where ***λ****_p,·_* = (*λ_p_*_1_*, …, λ_pk_*). Assuming the covariates are all on compatible scales, e.g., standardised continuous predictors, then ||***λ****_p,·_* || provides an effect size that is comparable across predictors; see the loading matrix and the construction of biplots in Hui et al. (2015) and van der Veen et al. (2023) for an analogous construction in the context latent variable models. Formal inference on the total effect of a covariate is also possible through, say, a multivariate Wald test of *H*_0_ : ||***λ****_p,·_*|| = 0 against *H*_1_ : ||***λ****_p,·_* || / = 0. This statistic is invariant to rotation of the latent axes, and tests the predictor’s total effect rather than its effect on any particular dimension. Inference on covariates in the dispersion model (3) is also possible via an analogous Wald tests on each coefficient in ***κ***. In our simulations in Section 4, we find the subsequent *p*-values for the Wald tests on ***λ****_p,·_* to be anti-conservative at small *n*, but to converge to the nominal level as the number of samples increases. Future research could potentially investigate the use of resampling-based techniques to handle the small sample case (taking inspiration from say, Warton et al., 2017).

We conclude this section by highlighting that the RRGDM output distinguishes two quantities: (1) The dissimilarity implied by the distance between the marginal positions (centroid) of two samples in latent space, reflecting location effects only, and (2) the expected dissimilarity between two samples drawn around those centroids, reflecting both location effects and dispersion. We illustrate this distinction in our first case study (Section 5.1). When centroids are close in latent space, the expected dissimilarity exceeds the centroid dissimilarity, with the gap largest when centroids coincide, as the expected dissimilarity is then governed entirely by dispersion. Because RRGDM parametrises dispersion explicitly via equation (3), both quantities can be obtained from the fitted model. Under isotropic dispersion, the contribution of dispersion to the expected dissimilarity can be derived analytically, since the latent distance between two samples follows a scaled non-central chi distribution. More generally, the expected dissimilarity is more conveniently approximated by Monte Carlo simulation from the fitted model.

## 4 Simulation study

We performed a numerical study based on simulating data from a RRGDM with known parameter values to assess the recovery of covariate effects and site coordinates used for ordination, along with the calibration of the Wald tests on dispersion and location effects. In detail, we set *n ∈ {*30, 60, 90, 120, 150*}* as the number of samples, the dimension of the latent space to either *k* = 2 or *k* = 3, and examined *m* = 3 standardised covariates ***x****_i_* entering (2) and one different standardised covariate *w_i_* entering the dispersion model (3). Pairwise dissimilarities were obtained from the resulting latent distances under a binomial likelihood with negative log link and 100 ‘trials’ per pair. We discarded replicates with clear signs of non-convergence, e.g., with absolute gradient above 0.1 or non-positive definite Hessian matrices returned from TMB. This led to at most 12 simulated datasets discarded in any scenario.

We assess parameter recovery by redrawing covariate effects from a normal distribution *λ_pl_ ∼ N* (0, 0.8^2^) and dispersion slopes from a uniform distribution *κ ∼ U* (*−*0.4, 0.4) for a total of 500 replicates. Recovery was assessed on rotation-invariant summaries, including the predictor modulus||***λ****_p,·_||* and the Procrustes error. We also assessed the recovery of latent coordinates (***ξ****_i_*) by comparing the Procrustes error (Schönemann, 1966) of the recovered coordinates. To assess the consequence of fitting lower latent dimensions than those involved in the data-generation process, we included scenarios where the data was simulated from *k* = 3, while fixing the latent dimension of the fitted RRGDM to *k* = 2.

With the dimension of the latent space correctly specified, the root mean squared error (RMSE) of all parameters more than halved between *n* = 30 and *n* = 150, with an even greater improvement for parameters in the dispersion sub-model *κ* and *κ*_0_ (Fig. 2A; Figs. S1–S2). A similar improvement was observed for the overall ordination of samples in latent space (Fig. 2B; Fig. S5 for the misspecified case). As it is common for dispersion parameters estimated via (unrestricted) maximum likelihood estimation (see Maestrini et al., 2026, and references therein), the intercept *κ*_0_ showed appreciable bias at small sample sizes, with a bias of *−*0.08 units at *n* = 30, although this decreased to *−*0.01 at *n* = 150. This pattern was only observed for *κ*_0_, while other parameters including *κ* did not show any appreciable bias at any sample size. As expected, misspecifying the RRGDM by fitting fewer latent dimensions than required (i.e., when the true *k* = 3) increased both biases and RMSE (Figs. S3–S4), with the variability corresponding to the unmodelled dimension partly absorbed by the sample-level error *u_il_* as reflected in a positive bias in *κ*_0_ that grew with sample size (Fig. S3).

**Figure 2:**
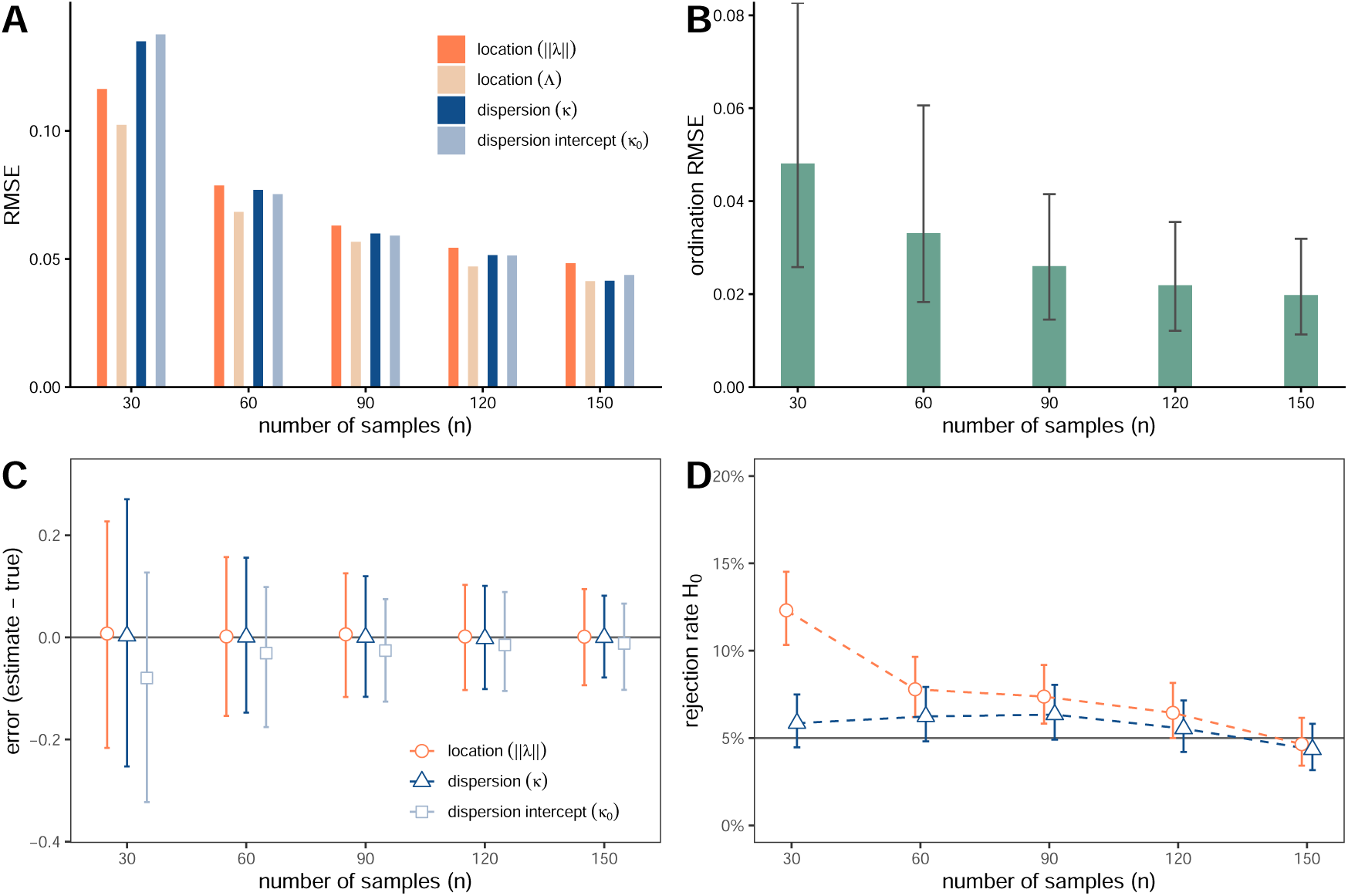
Wald-test calibration and parameter recovery from simulated data. (A) RMSE of RRGDM parameters. Location effects are assessed both through the individual modulus of each predictor (|| ***λ****_p,·_*||) and the Procrustes error of Λ. (B) RMSE (Procrustes error) of RRGDM ordination. Error bars show the spread of RMSE across replicates, delimiting the central 95% of values. (C) Bias, as represented by the average error (circle). Error bars show the spread of errors across replicates, delimiting the central 95% of simulated errors. (D) Calibration of Wald tests. Rejection rates represent the proportion of P-values lower than *α* = 0.05 under no simulated effect. Error bars represent the confidence intervals from a binomial distribution at 1000 replicates.

To assess the calibration of Wald tests, we fixed the covariate effect matrix Λ setting the third predictor to exactly zero, with both loadings freely estimated. The dispersion slope was likewise set to zero. We redrew random values for standardised covariates to obtain 1000 replicates. While the univariate Wald test for *κ* was close to nominal at all sample sizes (rejection rates [0.043, 0.063] at a nominal level of *α* = 0.05; Fig. 2D), the multivariate Wald test for the covariate effects *λ_p·_* was anti-conservative at small sample sizes, exhibiting a Type I error rate of 0.123 when *n* = 30. Type I error rates fell substantially with increasing sample size, with the distribution of *p*-values becoming consistent with a uniform distribution (Table S1; Fig. S6).

## 5 Case-studies

### 5.1 Drivers of compositional turnover in alpine plant communities

To illustrate the use of RRGDM in a realistic setting, we used the alpine plant community data published in Choler (2005) and available in the ade4 package (Dray & Dufour, 2007). This dataset contains counts of 82 plant species across 75 sites in the French Alps, spanning a pronounced gradient in snow-cover duration (Snow; indicating snow-melt day). We also included two recorded sources of disturbance: physical disturbance by periglacial processes (PhysD) and zoogenic disturbance by trampling and burrowing marmots (ZoogD), the latter coded as categorical predictors. Snow-cover duration and physical disturbance were scaled prior to fitting. We used Bray–Curtis dissimilarities with a binomial distribution, a negative log link function, and two latent variables. The two latent variables were sufficient to obtain a relatively low ordination stress, with only models improvement at *k* = 3 (Fig. S8), and without qualitative changes to model results (Table S2).

Model-based ordination of the fitted RRGDM in two dimensions showed a dominant gradient of variability in community composition due to snow-cover duration (Fig. 3A). Zoogenic disturbance also had a strong effect on community composition, approximately orthogonal to that of snow-cover duration. Both effects were statistically significant (Table 1, Fig. S7). Physical disturbance (PhysD) had a comparatively small but significant effect on community composition, whereas moderate zoogenic disturbance (ZoogDsome) was not distinguishable from baseline (no zoogenic disturbance). To represent sample-level error, marginal sample coordinates (determined only by fixed effects of environmental predictors) were compared against conditional coordinates (which include sample-level random effects) in the same ordination space (Fig. 3B).

**Figure 3:**
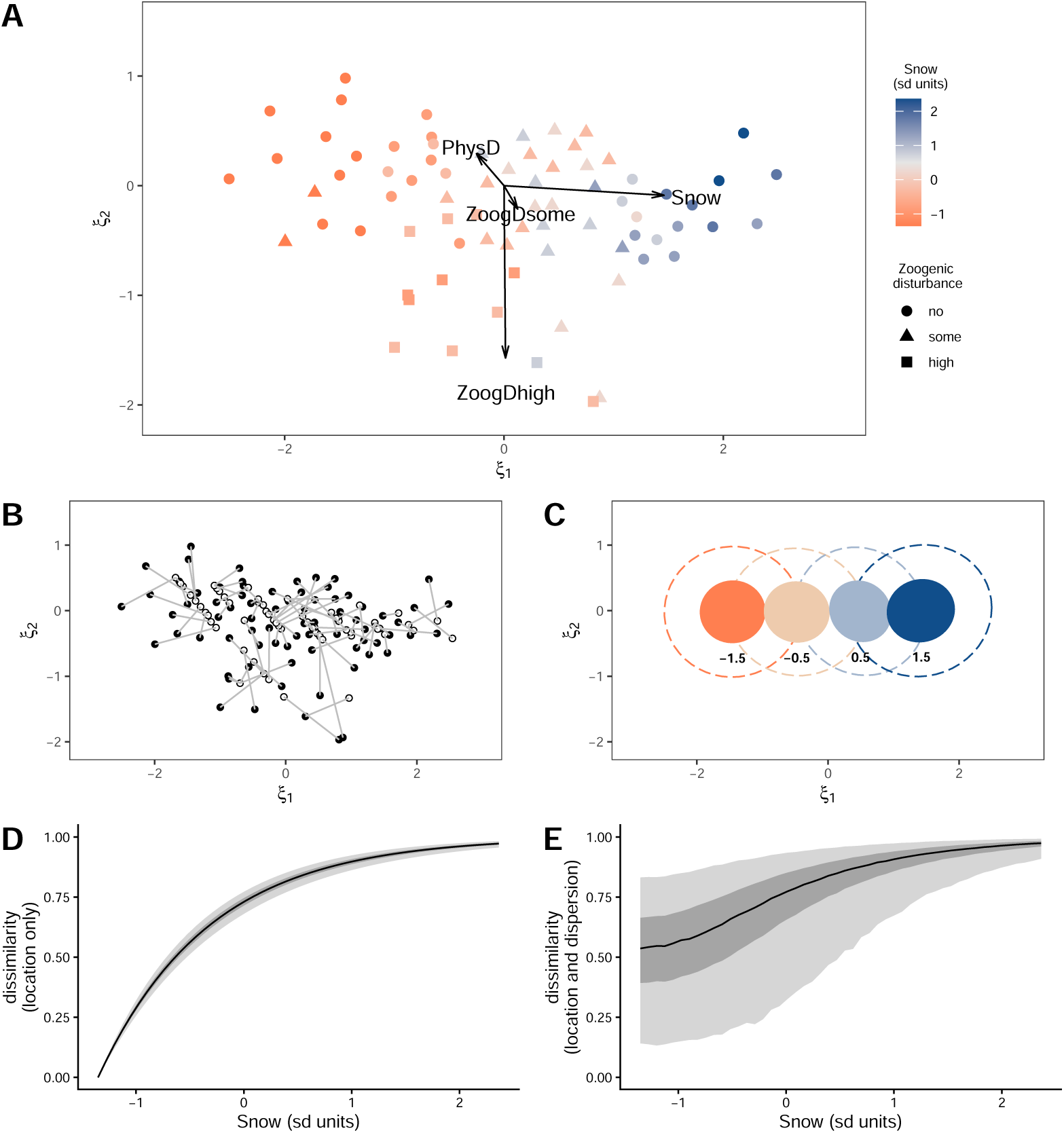
RRGDM ordination and expected dissimilarities (case study 1). (A) Biplot of the fitted model, showing the conditional community composition as captured by the two modelled latent axes, together with the environmental loadings of each predictor (arrows). Colours indicate the length of the snow-cover period (Snow); shapes indicate the degree of zoogenic disturbance (ZoogD). (B) Sample-level error, shown as the deviation of each sample from its marginal position (grey) to its conditional position (black). (C) Partial effect of Snow on latent community composition, shown for four values along the gradient (*±*1.5 standard deviation units). Filled and dashed circles indicate 50% and 95% of the expected sample density around each centroid. (D) Dissimilarity between average communities (centroids) along the snow-cover gradient, relative to the shortest period observed. The line is the median across parameter draws, and the shaded bands show 50% and 95% intervals reflecting parameter uncertainty. (E) Dissimilarity between samples along the same gradient, additionally accounting for the dispersion of samples around their centroids. Shaded bands show the central 50% and 95% of dissimilarities at the estimated parameter values. Panels D and E therefore reflect turnover in average composition (directional change) and dissimilarity between individual samples (including non-directional variability), respectively.

**Table 1:** Wald tests for environmental effects on (latent) composition of alpine plant communities. Significance codes: * *P <* 0.05, ** *P <* 0.01, *** *P <* 0.001.

| Variable | $\ \boldsymbol{\lambda}_{p,\cdot}\ $ | Wald $\chi^2$ | df | P-value |
| --- | --- | --- | --- | --- |
| Snow | 0.961 | 266.22 | 1 | <0.001*** |
| PhysD | 0.289 | 23.76 | 2 | <0.001*** |
| ZoogDsome | 0.154 | 1.67 | 2 | 0.433 |
| ZoogDhigh | 1.040 | 44.34 | 2 | <0.001*** |

The partial effect of snow-cover duration on latent community composition showed a near-complete turnover, with no overlap between the regions containing 95% of the expected sample density around the centroids at either end of the gradient (*±*1.5 SD) (Fig. 3C). Projected dissimilarities between average communities (centroids) likewise reflected a strong community turnover rising rapidly with distance along the gradient (Fig. 3D). Once the influence of compositional dispersion was included, expected Bray–Curtis dissimilarity between samples at equally short snow-cover periods exceeded 0.5 (Fig. 3E), indicating a substantial random variability in composition even under identical environmental conditions.

### 5.2 Combined productivity effects on average composition and dispersion

To illustrate the joint modelling of location and dispersion effects, we reanalysed the data from the mesocosm experiment in Chase (2010) where beta diversity among invertebrate communities in ponds with different trophic conditions was compared. The original analysis is a typical use-case for RRGDM because the hypothesis of interest concerns beta diversity both as changes in mean composition and compositional variability. Using a subset of the original data (McGlinn et al., 2020), consisting of 47 taxa across 30 mesocosms, we specified a binomial likelihood for Sørensen dissimilarities, which represent the proportion of unshared species between pairs of samples. Productivity (high or low) was included in both the mean (2) and dispersion models (3). We used *k* = 2 latent variables as stress and conditional fit (correlation-based *R*^2^) improved markedly from *k* = 1, but to a lesser extent with *k* = 3 (Fig. 4B).

**Figure 4:**
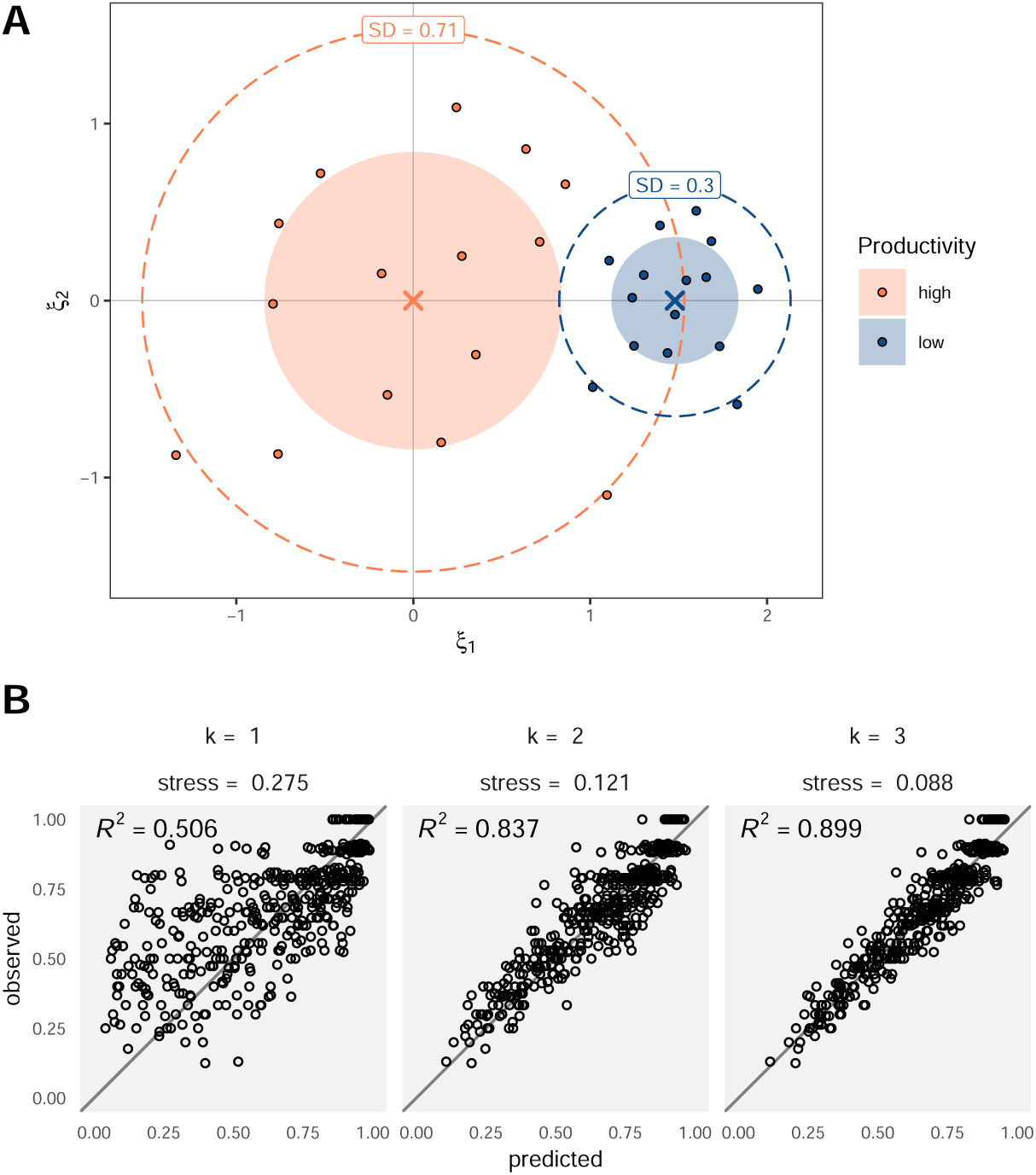
Effects of productivity on average composition and dispersion. (A) Fitted ordination showing conditional coordinates of samples (points), estimated group centroids (crosses), and the modelled dispersion of each group. Filled and dashed circles enclose 50% and 95% of the sample density around each centroid implied by the estimated dispersion *τ*^ (labelled values). (B) Observed against predicted dissimilarities for models with one, two, and three latent dimensions; grey lines indicate the 1:1 relationship. Correlation-based pseudo-*R*^2^ are also displayed.

The fitted model recovered the central result of the original study: productivity treatments differed in their dispersion (*z* = *−*4.42, *P <* 0.001), with highproductivity ponds showing considerably higher dispersion (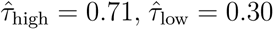; Fig. 4A). Additionally, it detected a difference in mean community composition between treatments (W*_df_*_=1_ = 51.36, *P <* 0.001). These two types of effect are conventionally assessed through separate non-parametric permutation procedures under which separating location and dispersion is a well-known difficulty (PERMANOVA and PERMDISP; Anderson, 2001; Anderson and Walsh, 2013; Warton et al., 2012), and patterns are represented within a separate ordination. Our RRGDM recovered effects on location and dispersion simultaneously, as well as latent dimensions that were used for ordination of samples and for the representation of environmental effects (productivity).

## 6 Discussion

In this article we propose RRGDM, a reduced-rank version of Generalised Dissimilarity Modelling (GDM) in which a low-dimensional latent space is used to capture compositional variability. A central contribution of our work is to resolve the mismatch between the pairwise scale at which beta diversity is measured, and the sample-level scale at which most processes driving community variability operate (Koleff et al., 2003; Socolar et al., 2016). Because RRGDM places environmental effects and residual variability on a common latent space, model-based ordination arises as a natural output. Additional structure such as categorical predictors, random effects, and predictors of dispersion can be seamlessly incorporated—in contrast to standard GDMs and other pairwise formulations. A simulation and two case studies demonstrate how RRGDM infers community turnover along environmental gradients and shifts in community dispersion.

In the low-dimensional latent space underlying RRGDM, sample-level errors displace each site from that determined by the measured covariates alone, and in doing so yields marginal and conditional coordinates of samples. RRGDM therefore shares similar motivation to spGDMM of White et al. (2024), which also imbues GDM with a likelihood and a flexible variance structure. The two models nonetheless differ in where the error structure is placed: spGDMM retains the pairwise mapping for environmental effects and captures residual sample-level variability through a separate latent dimension, whereas RRGDM specifies sample-level error directly on the latent space that carries the environmental effects. In the latter, systematic community turnover (i.e., location effects) and residual sample-level variability are therefore expressed in the same coordinates.

The capacity to formally model dispersion addresses a long-standing challenge in dissimilarity-based analyses: distinguishing shifts in mean composition from changes in compositional variability (Anderson et al., 2011; Warton et al., 2012). Because both processes influence average pairwise dissimilarities, the two signals can be conflated in standard permutation-based methods (Anderson, 2001; Anderson & Walsh, 2013), in models of community uniqueness (Tsang et al., 2023), and in conventional GDM (Ferrier et al., 2007; Hernández-Carrasco et al., 2026). RRGDM overcomes this by representing the two processes as separate components of a single model, with location effects determining a sample’s expected latent composition and dispersion effects determining the spread of samples around that position. We showcase this separation in our second case study, where a single model was used to estimate the difference in mean composition and dispersion between productivity treatments. Our approach is closest to that of Hernández-Carrasco et al. (2026), who capture the average contribution of community dispersion to observed dissimilarities as an additive, sample-level component within a pairwise formulation. Instead, RRGDM defines dispersion hierarchically as the variance of sample-level error terms, in direct analogy with distributional regression models developed for univariate responses (Merder et al., 2026; Stasinopoulos et al., 2018). This formal treatment of dispersion in dissimilarity models opens up exciting avenues for future research, from quantifying community homogenisation along invasion and stressor gradients (Olden & Rooney, 2006; Rolls et al., 2023), to assessing temporal and spatial changes in the predictability of community composition (Chase & Myers, 2011), and appropriately characterising the drivers of beta diversity.

Beyond separating ecological processes, the latent space architecture of RRGDMs provides the practical benefit of offering a model-based ordination as a direct output. Classical dissimilarity models return a scalar prediction for each pair of sites without a mapping of how samples are arranged relative to one another, meaning ordination often needs to be done in a separate, potentially *ad-hoc* downstream process for representation and visualisation (Chesters et al., 2019; Fitzpatrick et al., 2025; Lasram et al., 2015). By contrast, RRGDM allows sample coordinates and covariate effects—both on location and dispersion—to be visualised directly. For instance, our first case study showcases how partial effects of predictors and confidence intervals can be embedded within the RRGDM latent space, and thus visualised as an ordination. More advanced quantities such as community overlap can likewise be projected directly into this latent space, as demonstrated in our case studies (Fig. 4). Low-dimensional representations of this kind are fundamental to how multivariate data are explored and communicated in ecology (Hui et al., 2015; Legendre & Legendre, 2012; Legendre & Anderson, 1999; van der Veen et al., 2023), and thus RRGDM complements the existing ecological toolkit.

In parallel to dissimilarity methods, considerable progress has been made in modelling entire communities directly with joint species distribution models (JSDMs, Korhonen et al., 2025; Pollock et al., 2014; Tikhonov et al., 2020; Warton et al., 2015), from which beta diversity patterns can in principle be derived as a second step. Yet a broad class of applications concerns aggregate compositional change rather than the response of any particular taxon. Examples include testing treatment effects in experiments (where PERMANOVA and PERMDISP are currently the standard), comparing restored or potentially impacted communities to reference ones, quantifying biotic homogenisation (Olden & Rooney, 2006; Rolls et al., 2023), and comparing turnover rates across ecosystems and taxonomic groups (Nekola & White, 1999; Soininen et al., 2007). For these questions, dissimilarities are the quantities of direct interest. RRGDM models them directly with a small number of interpretable parameters, whereas a JSDM requires aggregating many species-specific responses after the fact. This brings two further practical advantages. First, the number of parameters in RRGDM does not depend on species richness, which makes it particularly convenient for very wide community matrices (e.g., microbial or macroinvertebrate surveys) where species identity is often secondary. Second, because the likelihood is fitted to the dissimilarities themselves, RRGDM favours the parameter values that best represent beta diversity, and may therefore capture it more reliably when it is driven by taxa that are hard to model individually, such as rare species or those with idiosyncratic responses. Yet, questions regarding certain species, trait–environment relationships (Niku et al., 2021; ter Braak, 2026), or predictions of species composition (Norberg et al., 2019) still require species-level models, and dissimilarity models are not a substitute for them. The two approaches are of course not mutually exclusive, and could be used in combination to obtain complementary information on the role of environmental drivers in structuring communities.

The main practical constraint on the implementation of RRGDMs is that the number of pairwise observations grows quadratically with the number of samples, an issue shared with all dissimilarity-based methods. For wide community matrices (i.e., many species and few sites), the approach remains efficient because computational cost is independent of species richness, but the computational cost might become limiting for surveys with many samples. Future research could explore limiting the number of neighbours per sample to reduce the scaling of pairs from quadratic to near-linear, a strategy that has been successfully applied in sparse implementations of multidimensional scaling and neighbour-graph embeddings based on dissimilarities (e.g., McInnes et al., 2020; Tenenbaum et al., 2000; van der Maaten & Hinton, 2008). Extensions in other directions follow from the model’s architecture. For instance, more flexible terms, such as splines and other smoothing terms for environmental effects, can be used to represent non-linear turnover along gradients (Fitzpatrick et al., 2013) and phenological changes in community composition (Lai, 2025), without altering the model formulation. Similarly, because sample-level errors are treated as random, they can be imbued with structured covariances so as to capture temporal and spatial dependencies among samples (Hui et al., 2023; Korhonen et al., 2025; Thorson et al., 2016). Such extensions are possible because RRGDM’s structure is closer to sample-level, making a range of existing solutions readily transferable.

We have demonstrated that modelling dissimilarities by placing samples in a shared latent space provides a suite of practical benefits to beta diversity analyses. Advancements include (i) the estimation of effects on average composition and on dispersion side by side in a single likelihood, (ii) handling of categorical predictors and random effects as in regular regression models, and (iii) the ability to visualise compositional variability and model parameters as an ordination. We expect RRGDM to be particularly useful where beta diversity itself is the focus, rather than the response of any single species, allowing researchers to detect, quantify, and visualise the sources of variability in community composition.

## Acknowledgements

G.K, is supported by Swiss National Foundation Project 10.006.102. F.K.C.H is supported by Australian Research Council Discovery Projects DP260100579 and DP240100143. J.D.T. is supported by a Rutherford Discovery Fellowship administered by the Royal Society Te Apārangi (RDF-18-UOC-007), and Te Pūnaha Matatini, a Centre of Research Excellence funded by the Tertiary Education Commission, New Zealand.

## Data Availability Statement

Data used in the two case studies are publicly available as part of existing software archived in The Comprehensive R Archive Network (CRAN; cran.r-project.org). An R package for fitting RRGDM is available on GitHub (github.com/dhercar/rrGDM), as well as the scripts used to produce the results in this manuscript (anonymous.4open.science/r/rrGDM_pub-76AE).

## Author Contributions

D.H.-C. and H.R.L. conceived the ideas; D.H.-C. and F.K.C.H. designed the methodology; D.H.-C. and G.K. developed the software; D.H.-C. and G.K. led the writing of the manuscript. All authors contributed critically to the drafts and gave final approval for publication.

## References

Allsup, C. M., George, I., & Lankau, R. A. (2023). Shifting microbial communities can enhance tree tolerance to changing climates. Science, 380(6647), 835– 840.

Anderson, M. J. (2001). A new method for non-parametric multivariate analysis of variance. Austral Ecology, 26(1), 32–46.

Anderson, M. J., Crist, T. O., Chase, J. M., Vellend, M., Inouye, B. D., Freestone, A. L., Sanders, N. J., Cornell, H. V., Comita, L. S., Davies, K. F., Harrison, S. P., Kraft, N. J. B., Stegen, J. C., & Swenson, N. G. (2011). Navigating the multiple meanings of *β* diversity: A roadmap for the practicing ecologist. Ecology Letters, 14(1), 19–28.

Anderson, M. J., & Walsh, D. C. I. (2013). PERMANOVA, ANOSIM, and the Mantel test in the face of heterogeneous dispersions: What null hypothesis are you testing? Ecological Monographs, 83(4), 557–574.

Barge, E. G., Leopold, D. R., Peay, K. G., Newcombe, G., & Busby, P. E. (2019). Differentiating spatial from environmental effects on foliar fungal communities of Populus trichocarpa. Journal of Biogeography, 46(9), 2001–2011.

Bray, J. R., & Curtis, J. T. (1957). An Ordination of the Upland Forest Communities of Southern Wisconsin. Ecological Monographs, 27 (4), 326–349.

Chase, J. M. (2010). Stochastic Community Assembly Causes Higher Biodiversity in More Productive Environments. Science, 328(5984), 1388–1391.

Chase, J. M., & Myers, J. A. (2011). Disentangling the importance of ecological niches from stochastic processes across scales. Philosophical Transactions of the Royal Society B: Biological Sciences, 366(1576), 2351–2363.

Chesters, D., Beckschäfer, P., Orr, M. C., Adamowicz, S. J., Chun, K.-P., & Zhu, C.-D. (2019). Climatic and vegetational drivers of insect beta diversity at the continental scale. Ecology and Evolution, 9(24), 13764–13775.

Choler, P. (2005). Consistent Shifts in Alpine Plant Traits along a Mesotopographical Gradient. Arctic, Antarctic, and Alpine Research, 37 (4), 444–453.

Claar, D. C., Tietjen, K. L., Cox, K. D., Gates, R. D., & Baum, J. K. (2020). Chronic disturbance modulates symbiont (Symbiodiniaceae) beta diversity on a coral reef. Scientific Reports, 10(1), 4492.

de Leeuw, J., & Mair, P. (2009). Multidimensional Scaling Using Majorization: SMACOF in R. Journal of Statistical Software, 31, 1–30.

Dias, F. S., Betancourt, M., Rodríguez-González, P. M., & Borda-de-Água, L. (2022). BetaBayes—A Bayesian Approach for Comparing Ecological Communities. Diversity, 14(10), 858.

Dice, L. R. (1945). Measures of the amount of ecologic association between species. Ecology, 26(3), 297–302.

Dray, S., & Dufour, A.-B. (2007). The ade4 Package: Implementing the Duality Diagram for Ecologists. Journal of Statistical Software, 22, 1–20.

Everitt, B. S., Landau, S., Leese, M., & Stahl, D. (2011, January). Cluster Analysis, 5th Edition. WILEY-BLACKWELL.

Ferrari, S., & Cribari-Neto, F. (2004). Beta Regression for Modelling Rates and Proportions. Journal of Applied Statistics, 31(7), 799–815.

Ferrier, S., Manion, G., Elith, J., & Richardson, K. (2007). Using generalized dissimilarity modelling to analyse and predict patterns of beta diversity in regional biodiversity assessment. Diversity and Distributions, 13(3), 252–264.

Fitzpatrick, M., Mokany, K., Manion, G., Nieto-Lugilde, D., Ferrier, S., Valavi, R., Lisk, M., Ware, C., Woolley, S., & Harwood, T. (2025, May). Gdm: Generalized Dissimilarity Modeling.

Fitzpatrick, M. C., Sanders, N. J., Normand, S., Svenning, J.-C., Ferrier, S., Gove, A. D., & Dunn, R. R. (2013). Environmental and historical imprints on beta diversity: Insights from variation in rates of species turnover along gradients. Proceedings of the Royal Society B: Biological Sciences, 280(1768), 20131201.

Fletcher, D., & Jowett, T. (2022). Single-fit bootstrapping: A simple alternative to the delta method. Methods in Ecology and Evolution, 13(6), 1358–1367.

Gower, J. C. (1966). Some Distance Properties of Latent Root and Vector Methods Used in Multivariate Analysis. Biometrika, 53(3/4), 325–338.

Griffiths, D. (1973). Maximum likelihood estimation for the beta-binomial distribution and an application to the household distribution of the total number of cases of a disease. Biometrics, 29, 637–648.

Hernández-Carrasco, D., Gillis, A. J., Lai, H. R., Siqueira, T., & Tonkin, J. D. (2026). Accounting for the Influence of Community Turnover Along Environmental Gradients on Compositional Uniqueness. Ecology Letters, 29(2), e70338.

Hui, F. K. C., Taskinen, S., Pledger, S., Foster, S. D., & Warton, D. I. (2015). Model-based approaches to unconstrained ordination. Methods in Ecology and Evolution, 6(4), 399–411.

Hui, F. K. C., Warton, D. I., Foster, S. D., & Haak, C. R. (2023). Spatiotemporal joint species distribution modelling: A basis function approach. Methods in Ecology and Evolution, 14, 2150–2164.

Jaccard, P. (1912). The distribution of the flora in the alpine zone. 1. New phytologist, 11(2), 37–50.

Knight, C. G., Nicolitch, O., Griffiths, R. I., Goodall, T., Jones, B., Weser, C., Langridge, H., Davison, J., Dellavalle, A., Eisenhauer, N., Gongalsky, K. B., Hector, A., Jardine, E., Kardol, P., Maestre, F. T., Schädler, M., Semchenko, M., Stevens, C., Tsiafouli, M. Α., … de Vries, F. T. (2024). Soil microbiomes show consistent and predictable responses to extreme events. Nature, 636(8043), 690–696.

Koleff, P., Gaston, K. J., & Lennon, J. J. (2003). Measuring beta diversity for presence–absence data. Journal of Animal Ecology, 72(3), 367–382.

Korhonen, P., Hui, F. K. C., Niku, J., Taskinen, S., & van der Veen, B. (2025). gllvm 2.0: fast fitting of advanced ordination methods and joint species distribution models. PeerJ, 13, e20338.

Kristensen, K., Nielsen, A., Berg, C. W., Skaug, H., & Bell, B. M. (2016). TMB: Automatic Differentiation and Laplace Approximation. Journal of Statistical Software, 70, 1–21.

Kruskal, J. B. (1964). Multidimensional scaling by optimizing goodness of fit to a nonmetric hypothesis. Psychometrika, 29(1), 1–27.

Lai, H. R. (2025). Model-based ordination for phenological studies: From controlling sampling bias to inferring temporal associations. Methods in Ecology and Evolution, 16, 1542–1546.

Lasram, F. B. R., Hattab, T., Halouani, G., Romdhane, M. S., & Loc’h, F. L. (2015). Modeling of Beta Diversity in Tunisian Waters: Predictions Using Generalized Dissimilarity Modeling and Bioregionalisation Using Fuzzy Clustering. PLOS ONE, 10(7), e0131728.

Legendre, P., & Legendre, L. (2012, July). Numerical Ecology. Elsevier.

Legendre, P., & Anderson, M. J. (1999). Distance-Based Redundancy Analysis: Testing Multispecies Responses in Multifactorial Ecological Experiments. Ecological Monographs, 69(1), 1–24.

Legendre, P., & De Caceres, M. (2013). Beta diversity as the variance of community data: Dissimilarity coefficients and partitioning. Ecology Letters, 16(8), 951– 963.

Lichstein, J. W. (2007). Multiple regression on distance matrices: A multivariate spatial analysis tool. Plant Ecology, 188(2), 117–131.

Maestrini, L., Hui, F. K. C., & Welsh, A. H. (2026). Restricted maximum likelihood estimation in generalized linear mixed models. Statistical Science, 41, 615– 636.

McGlinn, D. J., Xiao, X., Engel, T., & Oliver, C. (2020). mobr: Measurement of biodiversity in R (Version v2.0.0).

McInnes, L., Healy, J., & Melville, J. (2020, September). UMAP: Uniform Manifold Approximation and Projection for Dimension Reduction [arXiv:1802.03426 [stat.ML]].

Merder, J., Rigby, R. A., Mayr, A., Heller, G. Z., Kneib, T., Umlauf, N., De Bastiani, F., Stauffer, R., Tonkin, J. D., Logothetis, N., Zeileis, A., & Stasinopoulos, D. M. (2026). Distributional regression using generalized additive models for location, scale and shape. Nature Reviews Methods Primers, 6(1), 49.

Mokany, K., Ware, C., Woolley, S. N. C., Ferrier, S., & Fitzpatrick, M. C. (2022). A working guide to harnessing generalized dissimilarity modelling for biodiversity analysis and conservation assessment. Global Ecology and Biogeography, 31(4), 802–821.

Murtagh, F., & Legendre, P. (2014). Ward’s Hierarchical Agglomerative Clustering Method: Which Algorithms Implement Ward’s Criterion? Journal of Classification, 31(3), 274–295.

Nekola, J. C., & White, P. S. (1999). The distance decay of similarity in biogeography and ecology. Journal of Biogeography, 26(4), 867–878.

Niku, J., Brooks, W., Herliansyah, R., Hui, F. K. C., Taskinen, S., & Warton, D. I. (2019). Efficient estimation of generalized linear latent variable models. PloS one, 14, e0216129.

Niku, J., Hui, F. K. C., Taskinen, S., & Warton, D. I. (2021). Analyzing environmental-trait interactions in ecological communities with fourth-corner latent variable models. Environmetrics, 32, e2683.

Norberg, A., Abrego, N., Blanchet, F. G., Adler, F. R., Anderson, B. J., Anttila, J., Araújo, M. B., Dallas, T., Dunson, D., Elith, J., Foster, S. D., Fox, R., Franklin, J., Godsoe, W., Guisan, A., O’Hara, B., Hill, N. A., Holt, R. D., Hui, F. K. C., … Ovaskainen, O. (2019). A comprehensive evaluation of predictive performance of 33 species distribution models at species and community levels. Ecological Monographs, 89(3), e01370.

Olden, J. D., & Rooney, T. P. (2006). On defining and quantifying biotic homogenization. Global Ecology and Biogeography, 15(2), 113–120.

Peres-Neto, P. R., Legendre, P., Dray, S., & Borcard, D. (2006). Variation Partitioning of Species Data Matrices: Estimation and Comparison of Fractions. Ecology, 87 (10), 2614–2625.

Pollock, L. J., Tingley, R., Morris, W. K., Golding, N., O’Hara, R. B., Parris, K. M., Vesk, P. A., & McCarthy, M. A. (2014). Understanding co-occurrence by modelling species simultaneously with a Joint Species Distribution Model (JSDM). Methods in Ecology and Evolution, 5, 397–406.

Ramette, A. (2007). Multivariate analyses in microbial ecology. FEMS Microbiology Ecology, 62(2), 142–160.

Rolls, R. J., Deane, D. C., Johnson, S. E., Heino, J., Anderson, M. J., & Ellingsen, K. E. (2023). Biotic homogenisation and differentiation as directional change in beta diversity: Synthesising driver–response relationships to develop conceptual models across ecosystems. Biological Reviews, 98(4), 1388–1423.

Schönemann, P. H. (1966). A generalized solution of the orthogonal procrustes problem. Psychometrika, 31(1), 1–10.

Socolar, J. B., Gilroy, J. J., Kunin, W. E., & Edwards, D. P. (2016). How Should Beta-Diversity Inform Biodiversity Conservation? Trends in Ecology & Evolution, 31(1), 67–80.

Soininen, J., McDonald, R., & Hillebrand, H. (2007). The distance decay of similarity in ecological communities. Ecography, 30(1), 3–12.

Stasinopoulos, M. D., Rigby, R. A., & Bastiani, F. D. (2018). GAMLSS: A distributional regression approach. Statistical Modelling, 18(3-4), 248–273.

Tenenbaum, J. B., de Silva, V., & Langford, J. C. (2000). A Global Geometric Framework for Nonlinear Dimensionality Reduction. Science, 290(5500), 2319–2323.

ter Braak, C. J. (2026). Fourth-corner latent variable models overstate confidence in trait–environment relationships and what to use instead. Environmental and Ecological Statistics, 33, 77–96.

Thorson, J. T., Ianelli, J. N., Larsen, E. A., Ries, L., Scheuerell, M. D., Szuwalski, C., & Zipkin, E. F. (2016). Joint dynamic species distribution models: a tool for community ordination and spatio-temporal monitoring. Global Ecology and Biogeography, 25, 1144–1158.

Tikhonov, G., Opedal, Ø. H., Abrego, N., Lehikoinen, A., De Jonge, M. M., Oksanen, J., & Ovaskainen, O. (2020). Joint species distribution modelling with the R-package Hmsc. Methods in ecology and evolution, 11, 442–447.

Tsang, T. P. N., Bonebrake, T. C., Ponisio, L. C., & Cadotte, M. W. (2023). Controlling for the effects of environmental availability when testing how the environment determines community compositional uniqueness. Methods in Ecology and Evolution, 14(12), 3112–3122.

van der Maaten, L., & Hinton, G. (2008). Visualizing Data using t-SNE. Journal of Machine Learning Research, 9(86), 2579–2605.

van der Veen, B., Hui, F. K. C., Hovstad, K. A., & O’Hara, R. B. (2023). Concurrent ordination: Simultaneous unconstrained and constrained latent variable modelling. Methods in Ecology and Evolution, 14(2), 683–695.

van der Veen, B., Hui, F. K. C., Hovstad, K. A., Solbu, E. B., & O’Hara, R. B. (2021). Model-based ordination for species with unequal niche widths. Methods in Ecology and Evolution, 12, 1288–1300.

Warton, D. I., Blanchet, F. G., O’Hara, R. B., Ovaskainen, O., Taskinen, S., Walker, S. C., & Hui, F. K. C. (2015). So Many Variables: Joint Modeling in Community Ecology. Trends in Ecology & Evolution, 30(12), 766–779.

Warton, D. I., Thibaut, L., & Wang, Y. A. (2017). The PIT-trap – A “modelfree” bootstrap procedure for inference about regression models with discrete, multivariate responses. PloS one, 12, e0181790.

Warton, D. I., Wright, S. T., & Wang, Y. (2012). Distance-based multivariate analyses confound location and dispersion effects. Methods in Ecology and Evolution, 3(1), 89–101.

White, P. A., Frye, H. A., Slingsby, J. A., Silander, J. A., & Gelfand, A. E. (2024). Generative spatial generalized dissimilarity mixed modelling (spGDMM): An enhanced approach to modelling beta diversity. Methods in Ecology and Evolution, 15(1), 214–226.

Woolley, S. N., Foster, S. D., O’Hara, T. D., Wintle, B. A., & Dunstan, P. K. (2017). Characterising uncertainty in generalised dissimilarity models. Methods in Ecology and Evolution, 8(8), 985–995.

