## Supporting Information for "Reduced-Rank Generalised Dissimilarity Modelling (RRGDM): Model-Based Ordination of Dissimilarity Data"

#### S1 Separation of location and dispersion effects

We now show how effects on location and on dispersion enter the expected latent distance between samples. The latent position of sample $i$ in eq. (2) can be written compactly as

$$\boldsymbol{\xi}_i = \boldsymbol{x}_i \boldsymbol{\Lambda} + \boldsymbol{u}_i, \qquad \boldsymbol{u}_i \sim \mathcal{N}_k(\boldsymbol{0}_k, \tau_i^2 \boldsymbol{I}_k), \tag{S1}$$

where $\boldsymbol{\Lambda}$ is the $m \times k$ matrix of predictor loadings. Expanding the squared latent distance gives

$$\|\boldsymbol{\xi}_i - \boldsymbol{\xi}_j\|^2 = \underbrace{\Delta\boldsymbol{x}_{ij} \boldsymbol{\Lambda}\boldsymbol{\Lambda}^\top \Delta\boldsymbol{x}_{ij}^\top}_{\text{location}} + \underbrace{\|\boldsymbol{u}_i - \boldsymbol{u}_j\|^2}_{\text{dispersion}} + 2\,\Delta\boldsymbol{x}_{ij}\boldsymbol{\Lambda}(\boldsymbol{u}_i - \boldsymbol{u}_j)^\top, \tag{S2}$$

where $\Delta\boldsymbol{x}_{ij} = \boldsymbol{x}_i - \boldsymbol{x}_j$. Because $\boldsymbol{u}_i$ and $\boldsymbol{u}_j$ are independent and mean-zero, the expected cross term vanishes, and $\text{Var}(\boldsymbol{u}_i - \boldsymbol{u}_j) = (\tau_i^2 + \tau_j^2)\boldsymbol{I}_k$, so that

$$\mathbb{E}_{\boldsymbol{u}}\left[\|\boldsymbol{\xi}_i - \boldsymbol{\xi}_j\|^2\right] = \Delta\boldsymbol{x}_{ij} \boldsymbol{\Lambda}\boldsymbol{\Lambda}^\top \Delta\boldsymbol{x}_{ij}^\top \; + \; k\left(\tau_i^2 + \tau_j^2\right). \tag{S3}$$

This expression shows that the expected squared latent distance between two samples decomposes additively into a term governed entirely by location effects (through $\boldsymbol{\Lambda}$) and a term governed entirely by dispersion effects (through $\tau_i, \tau_j$, and hence $\boldsymbol{\kappa}$). Dispersion effects contribute an additive offset determined by each sample's own spread, whereas location effects depend on how far apart samples sit along the gradients.

Two properties of (S3) are worth noting. First, the location term depends on $\boldsymbol{\Lambda}$ only through $\boldsymbol{\Lambda}\boldsymbol{\Lambda}^\top$, which is invariant to orthogonal transformations (see main text). Second, the separation in (S3) is exact on the scale of squared latent distance only. Neither $\mathbb{E}_{\boldsymbol{u}}[\|\boldsymbol{\xi}_i - \boldsymbol{\xi}_j\|]$ nor $\mathbb{E}_{\boldsymbol{u}}[g^{-1}(\|\boldsymbol{\xi}_i - \boldsymbol{\xi}_j\|)]$ decompose additively, because the distance is a square root of (S3) and $g^{-1}$ is non-linear. The two contributions are therefore entangled in observed dissimilarities, which is what makes them difficult to separate in pairwise analyses. One of RRGDM's main contributions is resolving this by estimating $\boldsymbol{\Lambda}$ and $\boldsymbol{\kappa}$ as distinct components of a single likelihood.

#### S2 Initialisation strategy for optimisation

To reduce computation time and assist in model convergence, starting parameter values are obtained with the following strategy:

1. We apply Principal Coordinates Analysis (Gower, 1966) to the link-transformed dissimilarity matrix to obtain initial sample scores $\boldsymbol{\xi}_i$ for the latent variables.

2. We then perform a distance-based Redundancy Analysis (Legendre & Anderson, 1999), extracting out environmental covariate effects $\boldsymbol{\lambda}_l$ and site residuals $u_{il}$ for the top $k$ constrained axes.

3. To obtain initial values for covariate effects that satisfy the aforementioned identifiability constraints, we rotate the $\boldsymbol{\Lambda}$ from step 2 into a lower-triangular form with positive diagonal via a QR decomposition, and apply the same rotation to the residual site scores to obtain a final set of initial values for $\boldsymbol{U}$.

4. Finally, dispersion slopes $\boldsymbol{\kappa}$ are initialised at zero, and $\kappa_0$ at the logarithm of the pooled standard deviation of the entries of $\boldsymbol{U}$.

This procedure improves efficiency and numerical stability without affecting the model formulation. Note the above starting parameter values may also be jittered several times (analogous to Niku et al., 2019, for latent variable models), if multiple restarts are utilized.

### S3 Simulation study: Supplementary tables and figures

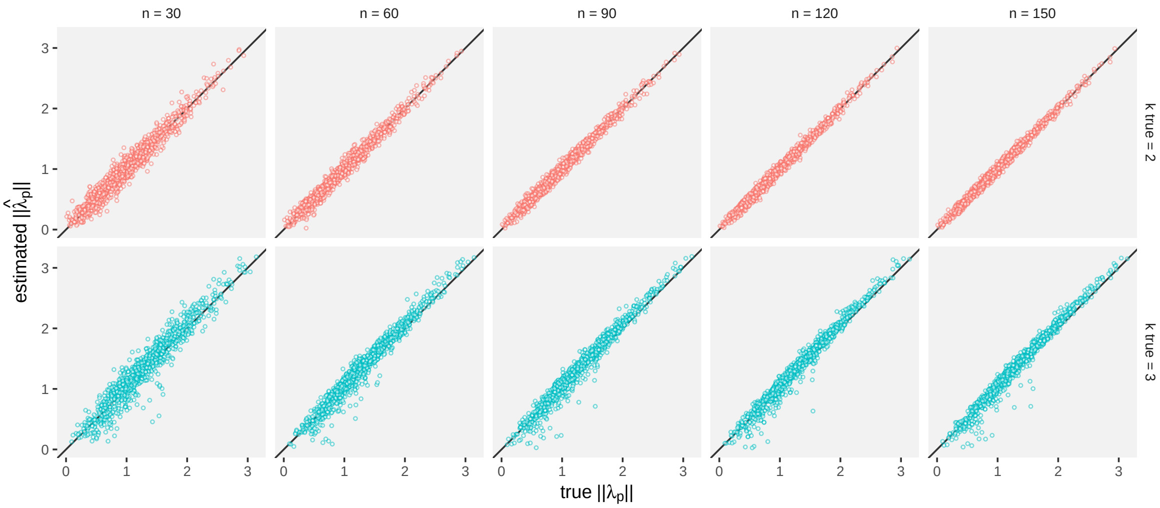


Figure S1: Estimated versus simulated (true) location effects ($\|\boldsymbol{\lambda_p}\|$). Top panels ($k = 2$) were fitted with the same number of dimensions as the data-generation process, whereas bottom panels were purposefully misspecified with one more dimension ($k = 3$) in the data-generation process.

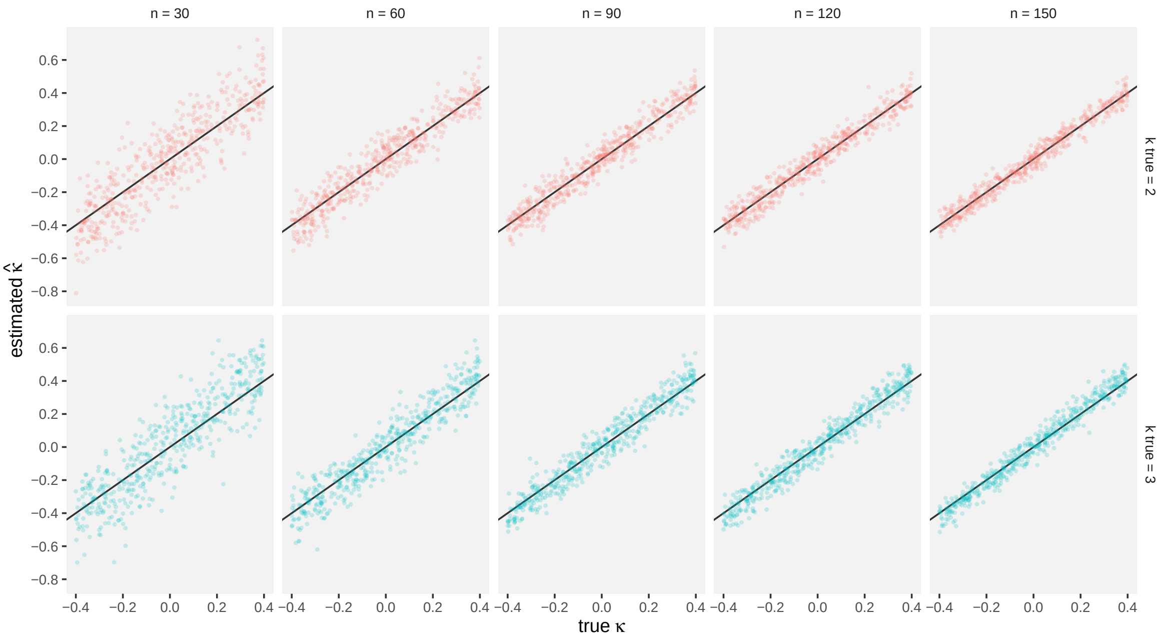


Figure S2: Estimated versus simulated (true) dispersion effects ($\hat{\kappa}$). Top panels ($k = 2$) were fitted with the same number of dimensions as the data-generation process, whereas bottom panels were purposefully misspecified with one more dimension ($k = 3$) in the data-generation process.

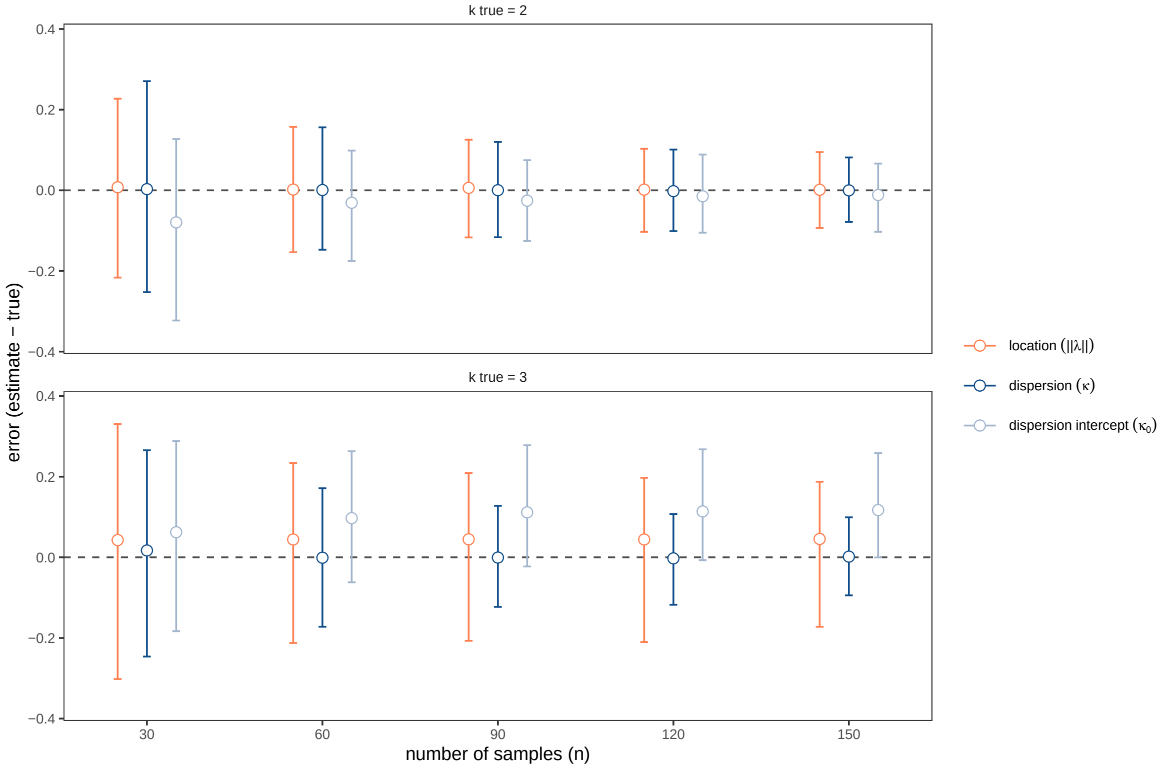


Figure S3: Estimation bias, as represented by the average error across replicates (circle). Error bars show the spread of errors across replicates, delimiting the central 95% of simulated errors. Replicates in the top panel ($k = 2$) were fitted with the same number of dimensions as the data-generation process, whereas replicates in the bottom panel were purposefully misspecified with one more dimension ($k = 3$) in the data-generation process.

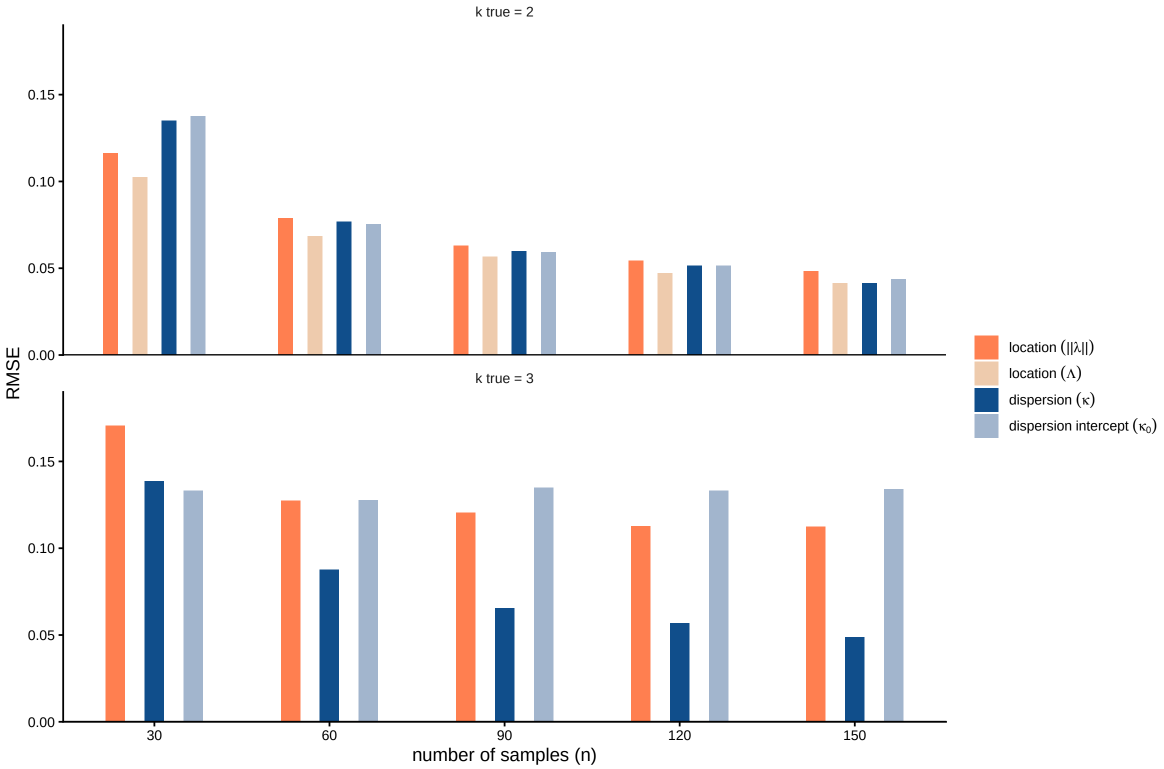


Figure S4: Error (RMSE) of fitted parameters for replicates where the true dimensions matched those in the fitted model ($k = 2$) or where dimensions were misspecified by estimating one less dimension than the data-generation process ($k = 3$). We compared the modulus of environmental loadings ($\|\lambda_p\|$) and the Procrustes error ($\Lambda$) between true and recovered loading matrices $\Lambda$. The latter was omitted for scenarios with misspecified dimensionality.

Table S1: Calibration of the Wald test. $c_{wald}$ is the mean Wald $\chi^2$ statistic across replicates divided by its expected value under the null hypothesis (its degrees of freedom), so values above 1 indicate inflated test statistics (anti-conservative P-values) and values below 1 indicate conservative tests. For the set of replicates in each scenario, we also compare the resulting distribution of P-values to the uniform distribution $\mathcal{U}(0, 1)$ with a Kolmogorov–Smirnov test ($P_{\text{K-S}}$). Significance codes: * $P < 0.05$, ** $P < 0.01$, *** $P < 0.001$.

| Parameter | Sample size | Usable replicates | $c_{\text{wald}}$ $\bar{\chi}^2$ | $P_{\text{K-S}}$ |
|---|---|---|---|---|
| Location | 30 | 991 | 1.38 | <0.001*** |
| Location | 60 | 988 | 1.14 | 0.002** |
| Location | 90 | 990 | 1.11 | 0.047* |
| Location | 120 | 992 | 1.05 | 0.507 |
| Location | 150 | 989 | 1.01 | 0.808 |
| Dispersion | 30 | 991 | 1.09 | 0.395 |
| Dispersion | 60 | 994 | 1.12 | 0.081 |
| Dispersion | 90 | 993 | 1.08 | 0.490 |
| Dispersion | 120 | 993 | 1.04 | 0.844 |
| Dispersion | 150 | 989 | 0.98 | 0.982 |

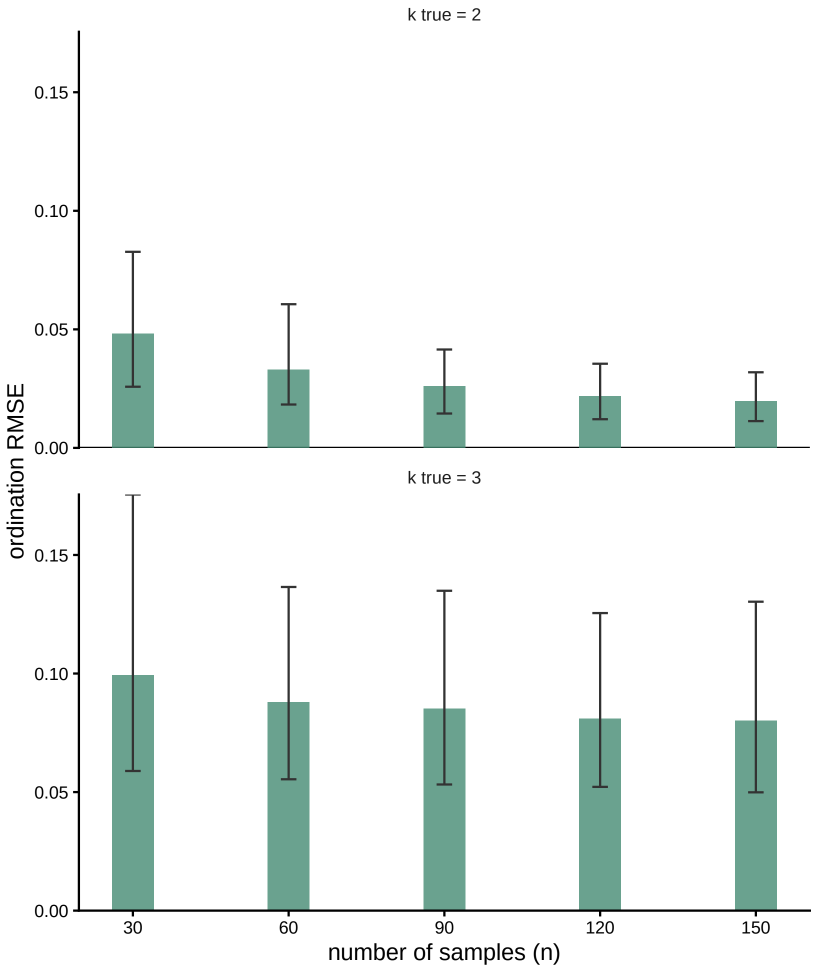


Figure S5: Procrustes error (RMSE) of estimated latent coordinates ($\hat{\boldsymbol{\xi}}_i$) for replicates where the true dimensions matched those in the fitted model ($k = 2$) or where dimensions were misspecified by estimating one less dimension than the data-generation process ($k = 3$). Error bars show the spread of RMSE across replicates, delimiting the central 95% of values.

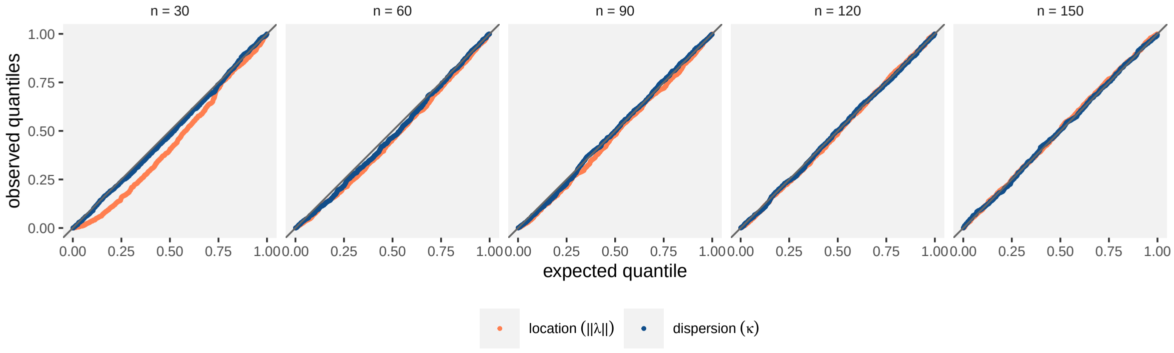


Figure S6: Comparison of expected versus observed quantiles (QQ-plot) of P-values obtained from the Wald test.

### S4 Case studies: Supplementary tables and figures

Table S2: Wald tests for environmental effects on (latent) composition of alpine plant communities. RRGDM fitted with $k = 3$ latent dimensions. Significance codes: * $P < 0.05$, ** $P < 0.01$, *** $P < 0.001$.

| Variable | $\|\boldsymbol{\lambda}_{p,\cdot}\|$ | Wald $\chi^2$ | df | P-value |
|---|---|---|---|---|
| Snow | 0.934 | 347.807 | 1 | <0.001*** |
| PhysD | 0.305 | 36.640 | 2 | <0.001*** |
| ZoogDsome | 0.246 | 5.679 | 3 | 0.128 |
| ZoogDhigh | 0.817 | 37.086 | 3 | <0.001*** |

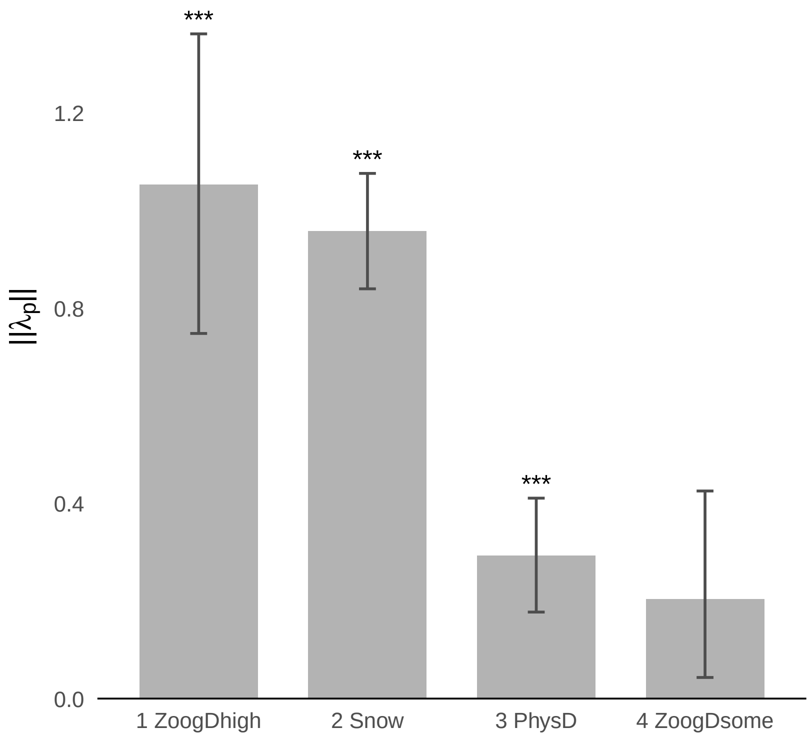


Figure S7: Effect sizes of environmental predictors on latent composition (case study 1), from an RRGDM fitted with $k = 2$ latent dimensions. Error bars show 95% confidence intervals obtained from 1000 draws from the asymptotic distribution of parameter estimates. Significance codes indicate the result of a multivariate Wald test: * $P < 0.05$, ** $P < 0.01$, *** $P < 0.001$.

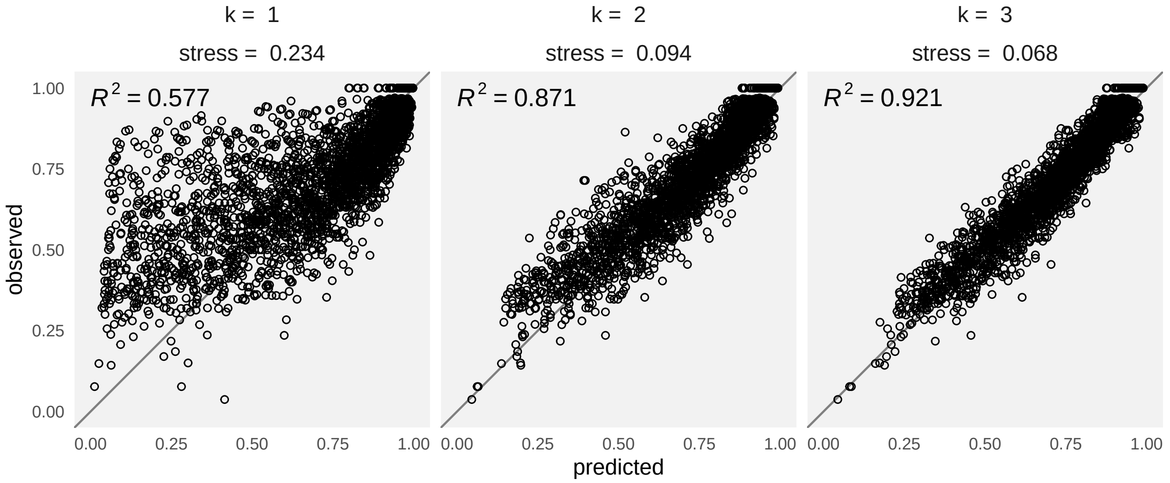


Figure S8: Observed against predicted dissimilarities (case study 1). Panels represent RRGDMs fitted with one ($k = 1$), two ($k = 2$), and three latent dimensions ($k = 3$). Grey lines indicate the 1:1 relationship.

### References

Gower, J. C. (1966). Some Distance Properties of Latent Root and Vector Methods Used in Multivariate Analysis. *Biometrika*, *53*(3/4), 325–338.

Legendre, P., & Anderson, M. J. (1999). Distance-Based Redundancy Analysis: Testing Multispecies Responses in Multifactorial Ecological Experiments. *Ecological Monographs*, *69*(1), 1–24.

Niku, J., Brooks, W., Herliansyah, R., Hui, F. K. C., Taskinen, S., & Warton, D. I. (2019). Efficient estimation of generalized linear latent variable models. *PloS one*, *14*, e0216129.
